# Targeting autoimmunity in Rheumatoid Arthritis with nanoparticles displaying Liprin-1 peptide

**DOI:** 10.64898/2026.01.09.698578

**Authors:** Anthony Rosa, Alessia Raneri, Michela Campolo, Emanuela Esposito, Elisa Gecchele, John Edward Butler, Julian K-C. Ma, Nidhi Sofat, Santiago Figueira, Denise Pivotto, Alison McCormick, Elena Bartoloni, Roberta Zampieri, Linda Avesani

**Affiliations:** Diamante SB srl, Viale del Lavoro, 33. 37135 Verona (Italy); Dipartimento Biotecnologie, Università di Verona, Strada Le Grazie 15. 37134 Verona (Italy); Department of Chemical, Biological, Pharmaceutical and Environmental Sciences; University of Messina. Viale Ferdinando Stagno D’Alcontres, 31- 98166 Messina (Italy); Institute for Infection and Immunity, School of Health and Medical Sciences, City St. George’s University of London, Cranmer Terrace, London SW17 0RE (UK); Palobiofarma S.L., Carretera Pamplona 1 31191 Galar (Spain); Department of Chemical Engineering, University of California, 3001 Ghausi Hall, Davis, CA (USA); Department of Internal Medicine, University of Perugia (Italy)

## Abstract

Nanoparticle-based strategies offer a promising tool for inducing antigen-specific immune tolerance across various autoimmune conditions, by acting as master switch to turn-off the autoimmune response. Building on our previous work demonstrating the therapeutic potential of plant-made nanoparticles in rheumatoid arthritis, we present a systematic evaluation of key parameters—including dosing regimen, route of administration, and immunization schedule—to optimize both efficacy and safety. We developed virus-based nanomaterials expressing Tomato Bushy Stunt Virus (TBSV) nanoparticles engineered to display the immunodominant Liprin-1 peptide in the *Nicotiana benthamiana* plant platform. The mechanisms responsible for the observed protective effects against rheumatoid arthritis were also investigated.

Our findings highlight the critical role of repeated intravenous administration and precise dosing in promoting regulatory T cell (Treg) induction and cytokine modulation. Furthermore, we dissected the individual contributions of the Liprin-1 peptide and the viral capsid scaffold in driving immune tolerance.

These results support the potential of plant-derived nanoparticles as a versatile and effective platform for antigen-specific immunotherapy, with rheumatoid arthritis serving as a proof-of-concept model for broader applications in the field of autoimmunity.

**One Sentence Summary:** Plant-made nanomaterials induce tolerance in arthritis models via repeated IV dosing, triggering an antigen-specific regulatory immune response.

## INTRODUCTION

Rheumatoid arthritis (RA) is a chronic, progressive and debilitating autoimmune disease characterized by the malfunction of various immune cell types and signaling network, resulting in an improper tissue repair process that primarily damages the joints but also affects the lungs and the vascular system^1^. In 2020, an estimated 17.6 million people worldwide had RA and this number is projected to increase to 31.7 million by 2050 based on forecasted change in population age^2^.

Since the first treatments for symptomatic relief (non-steroidal anti-inflammatory drug and glucocorticoids), RA treatment has evolved to include disease modifying anti-rheumatic drugs (DMARD), ranging from methotrexate to advanced biological and targeted synthetic DMARDs, yet these treatments still cannot fully address the underlying immune disorders, requiring lifelong use with the possibility of severe adverse effects^3^.

For decades, numerous studies for autoimmune diseases therapy, have aimed to restore tolerance, which is the ability of the immune system to ignore self-antigens while appropriately attacking foreign ones^4^. Tolerance induction may be achieved either by 1) anergy or deletion of specific-autoreactive T-cells or by 2) converting autoreactive T cells to a Treg phenotype^5^. The ultimate goal of the latter approach is to re-program autoreactivity to induce bystander immunoregulation by the capacity of one or a few autoreactive regulatory T- and/or B-cell specificities, which will ultimately blunt the activation of non-cognate effector lymphocytes. A major advantage of this “active” approach, defined as Antigen-Specific Immunotherapy (ASI), is that sustained activation and expansion of regulatory cells targeting specific epitopes should be able to block the recruitment and activation of all other adaptive and innate immune cells that take part in the disease process^6^.

Several products that have shown promising results are now at various stages of the clinical development pipeline^3^; among these, nanoparticles hold significant promise for development, due to a better understanding of their mechanisms of action in different autoimmune diseases settings^7^. In tolerance induction, nanoparticles using inorganic and/or organic materials can act as vehicles for drug and/or antigen delivery to phagocytes, as well as direct T-cell targeting compounds^8^. By specific design, the structural properties of nanomaterials can contribute to the pharmacokinetic and pharmacodynamics of a target compound, becoming an essential ingredient of the final product rather than an inert component^6^.

We previously demonstrated in preclinical studies that plant-made Virus Nano-Particles (VNPs) can be successfully used to induce tolerance both for autoimmune diabetes prevention and for RA treatment^9^; VNPs, derived from two distinct plant viruses, were genetically engineered to display immunodominant peptides related to the target autoimmune diseases on their outer surface. They were then propagated in *Nicotiana benthamiana* plants through a rapid expression system^9^. Notably, commercial approval was granted for the first time in 2023 for a COVID-19 VLP vaccine produced using the same bio factory, *N.benthamiana*^10^. The workflow exploited enables safe production of high titers of monodispersed VNPs in short timeframes, thus paving the way for their future advancement as a therapy for tolerance induction in autoimmune diseases. We recently established a GMP-compliant facility for the manufacturing of clinical-grade VNPs meant to progress to a first in human (FIH) clinical study of the safety of the particles^11^.

Starting from this first proof-of-concept of the use of plant-made VNPs for tolerance induction, we further explore their therapeutic potential, namely, TBSV.pLip, in RA treatment by pre-clinical studies specifically designed to determine their pharmacological activity. Furthermore, we gathered evidence to define the mechanism of action of TBSV.pLip in a pre-clinical model.

## RESULTS

### Nanoparticles manufacturing in a TBSV/plant-based system

TBSV WT and TBSV.pLip virus nanoparticles (VNPs) were produced in *N. benthamiana* plants, following previously established protocols^9^ and a recently optimized production workflow^11^. For TBSV.pLip, the Liprin-1–derived peptide (pLip) coding sequence was genetically fused to the C-terminus of the viral coat protein gene within the TBSV genome, resulting in surface display of the peptide on the exterior of the assembled viral capsid. Thus, pLip is structurally incorporated into the VNP through genetic fusion and is presented on the particle external surface.

VNPs were purified from infected leaf tissue^9^ and quantified via Enzyme-Linked Immunosorbent Assay (ELISA)^11^. VNPs were then characterized by agarose gel electrophoresis and Dynamic Light Scattering (DLS) to assess integrity and size distribution (Figure 1A–B). DLS analysis detected no significant difference in hydrodynamic radius between TBSV.pLip and TBSV WT (Figure 1B). Extraction of nanoparticles occurred under controlled conditions and endotoxin levels was below the acceptable standard of 1.50 EU/ml^12^.

**Fig. 1.**
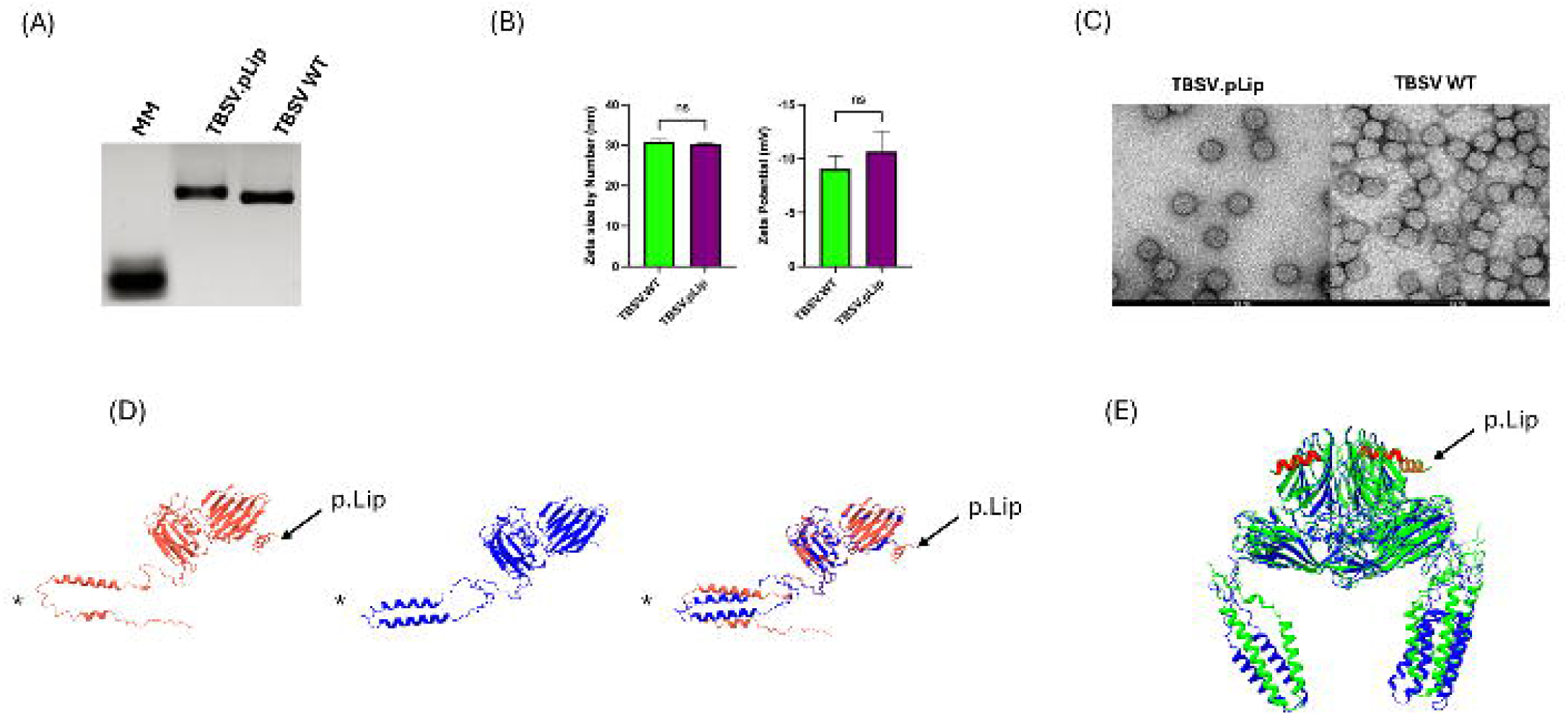
Virus nanoparticles characterization. (A) Agarose gel electrophoretic patterns of 1 µg of purified TBSV.pLip and TBSV WT. MM: molecular marker. (B) Characterization of TBSV.pLip and TBSV WT by DLS evaluation, measuring size distribution by number and their Zeta Potentials. The reported data represent the mean value of 50 measurements +/- standard deviation. The data groups were compared by Student T-test. n.s.: not significant. (C) TEM images of TBSV.pLip and TBSV WT. (D) Structural modeling of TBSV.pLip and TBSV WT coat protein monomers. The sequence alignment shows a score of 1933.3. The root-mean-square deviation (RMSD) is 0.308 Å for 312 pruned atom pairs and 10.208 Å across all 388 pairs, indicating a mismatch in the unstructured C-terminal loops that fail to align (asterisk). In contrast, the surface-exposed β-sheet regions are almost identical, except for the Liprin-1 peptide helix (black arrows). (E) Structural modeling of TBSV.pLip and TBSV WT coat protein trimers.

Zeta potential measurements revealed that the surface charge of the VNPs is negative, showing an average of -10.67 mV for TBSV.pLip and -9.05 mV for TBSV WT (Figure 1B). Transmission electron microscopy (TEM) analysis confirmed that both formulations exhibited uniform spherical morphology, (Figure 1C). Modelling of the TBSV CP monomer indicates that the surface-exposed β-sheet motifs are highly conserved, except for the Liprin-1 peptide (pLip), which is sequestered within the interstitial regions at subunit interfaces (Figure 1D-E). These findings support experimental data (Figure 1B-C), indicating that TBSV.pLip particles, despite incorporation of the peptide, do not exhibit increased dimensions compared to TBSV WT.

### Administration routes for tolerance induction

We first sought to validate our previous observations obtained in female DBA/1J CIA mice treated with TBSV.pLip (seven intraperitoneal injections; n = 3 per group)^9^ by assessing tolerance induction in male DBA/1J CIA mice. Male animals were selected based on prior evidence indicating a more rapid disease onset and increased arthritis severity in this sex^13,14^, thus providing a more stringent context for therapeutic evaluation. Using a larger cohort (n = 10), this experiment was designed to confirm the robustness of the tolerogenic effect under more severe disease conditions.

In parallel, considering that the route of administration can critically influence immunological outcomes, yet is rarely systematically compared in tolerance-induction studies, we evaluated its impact on therapeutic efficacy in CIA mice within the same experimental setting. We had previously demonstrated that seven intraperitoneal (i.p.) injections administered five days apart were effective in treating arthritis female DBA/1J CIA induced^9^. Using the same dosing schedule (Figure 3A), we compared four administration routes: intraperitoneal (i.p.), subcutaneous (s.c.), intravenous (i.v.), and oral (p.o.) in male DBA/1J mice to determine their ability to halt disease progression (Figure 2A).

**Fig. 2.**
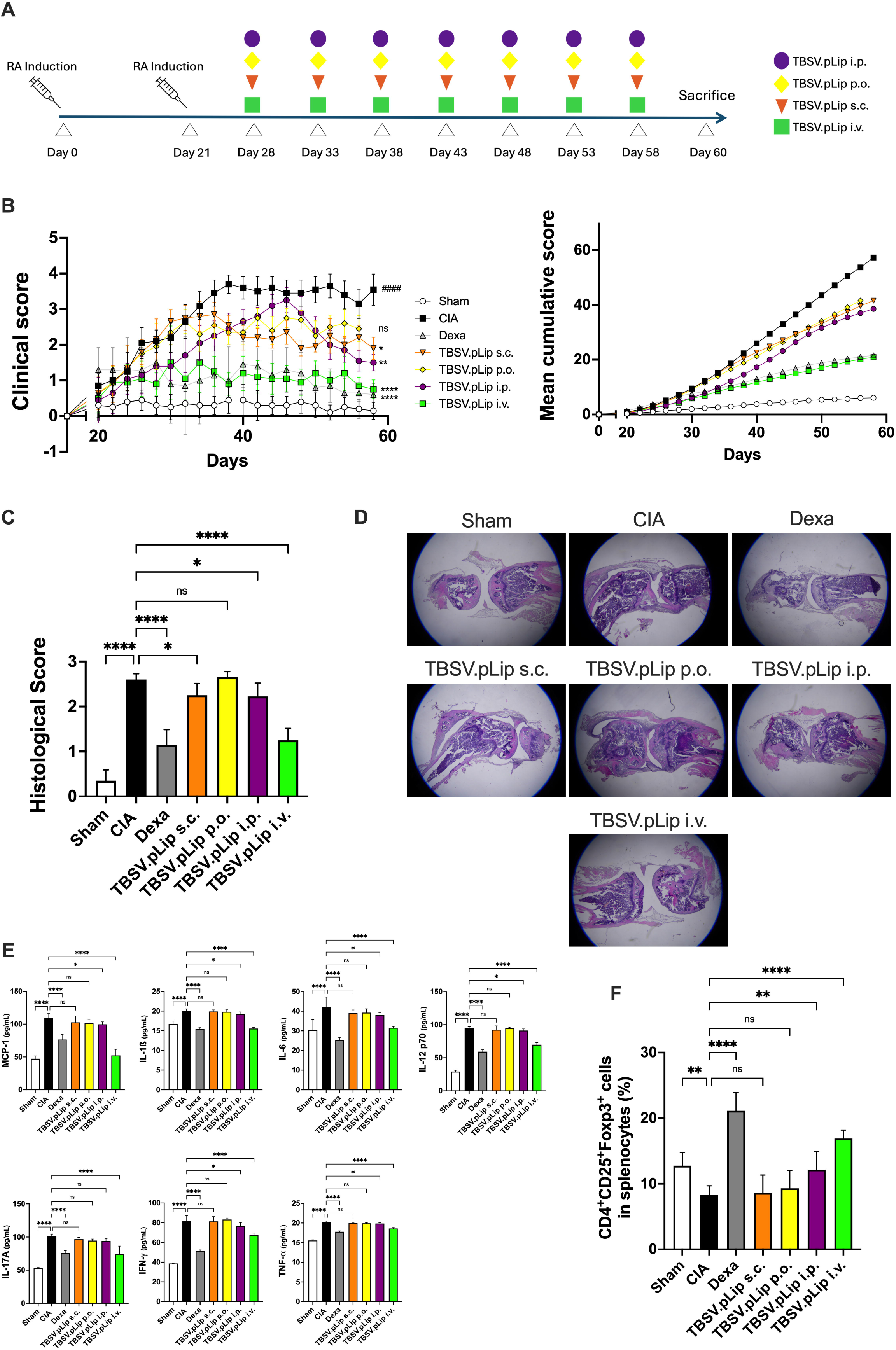
Impact of the route of administration on TBSV.pLip-mediated tolerance induction. (A) Outline of the experimental design. 8 weeks-old mice received on day 0 and boosted on day 21 intradermally at the basis of the tail 200 µg of collagen with Complete Freund’s adjuvant (CFA). The animals (n=10 for each group) were divided into 7 groups on day 28 based on clinical score and/or paw volume. 7 doses of 50µg of TBSV.pLip were administered by different injection routes: intra-peritoneal (i.p.), intra-venous (i.v.), sub-cutaneous (s.c.) and oral (p.o.); control groups were: untreated mice (CIA), sham group and a group receiving dexamethasone (1 mg/kg) every other day (Dexa). The animals were sacrificed 60 days after CIA induction. (B) *Left part:* Comparison of clinical score of all the animal groups. Clinical parameters such as slight edema and erythema limited to ankle, slight edema and erythema from the ankle to the tarsal bone, moderate edema and erythema the ankle to the tarsal bone, edema and erythema from the ankle to the entire leg have been considered for clinical score. Statistical significance was determined by 1-way ANOVA. *: p<0.05, **: p<0.01, ****: p<0.001 in comparison with the CIA group; ####: p<0.001 in comparison with the sham group, ns: not significant. *Right part:* corresponding cumulative scores. (C) Comparison of histological score in ankle-joints of analyzed animal groups. Statistical significance was determined by 1-way ANOVA: *: p<0.05; ****: p<0.001; ns: not significant in comparison with CIA. After 60 days, ankle-joints were evaluated by addressing clinical score from 1 to 4: (1) slight edema and erythema limited to ankle; (2) slight edema and erythema from the ankle to the tarsal bone; (3) moderate edema and erythema the ankle to the tarsal bone; (4): edema and erythema from the ankle to the entire leg. (D) Representative images of histological sections of ankle-joints of analyzed animal groups. (E) Pro-inflammatory cytokines profiles (MCP-1, IL-1β, IL-6, IL-12p70, IL-17A, INF-γ, TNF-α,) in sera of analyzed mice. Statistical significance was assessed by 1-way ANOVA: *: p<0.05, **: p<0.01; ***: p<0.005; ****: p<0.001 in comparison to CIA. (F) Percentage of CD4+CD25+FoxP3+ Treg cells among CD4+ cells in splenocytes of mice analyzed by flow cytometric analysis. Statistical significance was determined by 1-way ANOVA: *: p<0.05, **: p<0.01; ***: p<0.005; ****: p<0.001 in comparison to CIA.

**Fig. 3.**
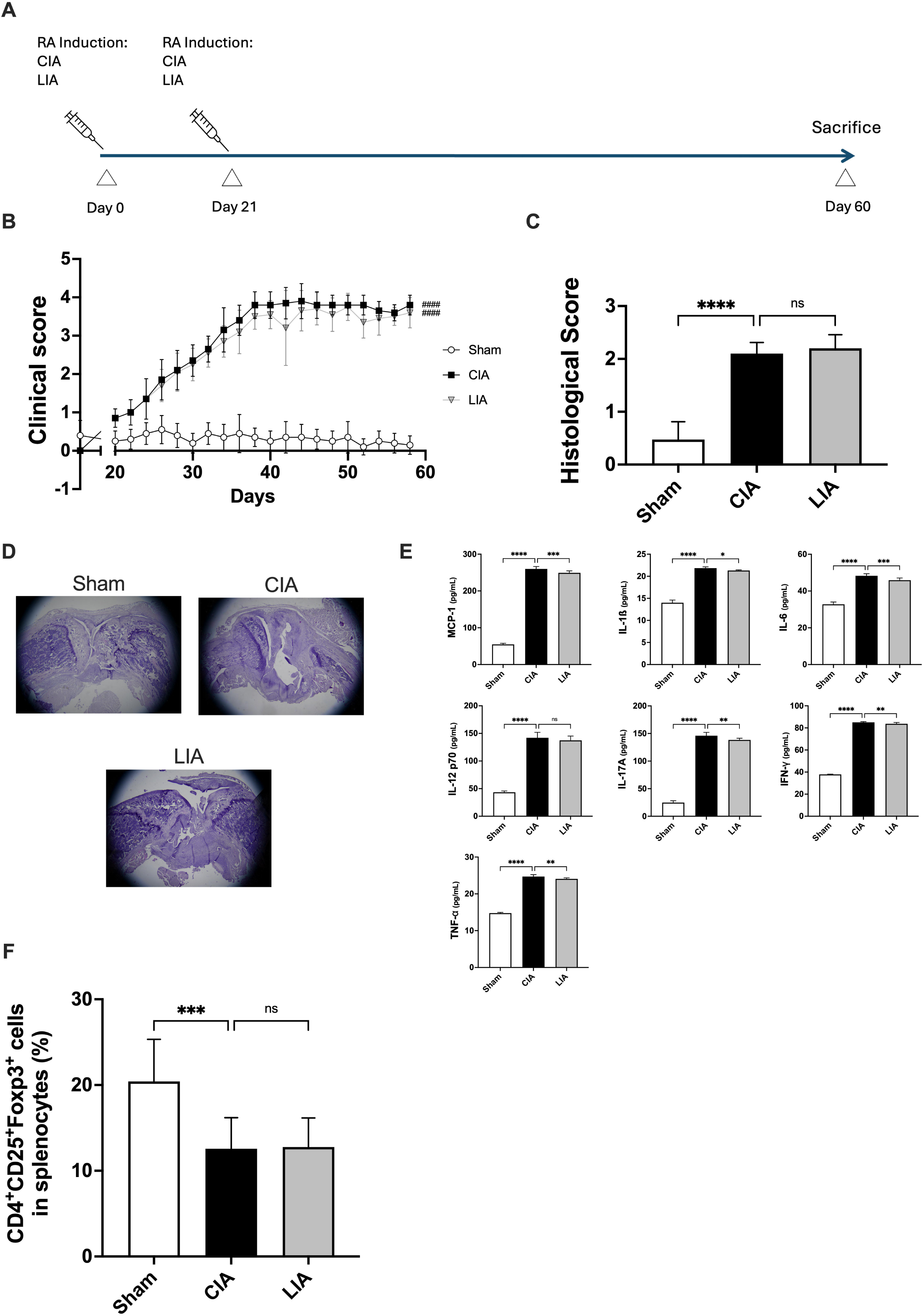
Arthritis induction with LIP1 peptide. (A) Outline of the experimental design. 8 weeks-old mice received on day 0 and boosted on day 21 intradermally at the basis of the tail 200 µg of collagen an Liprin-1 peptide both with Complete Freund’s adjuvant (CFA). The animals (n=10 for each group) were sacrificed 60 days after CIA and LIA induction. (B) *Left part:* comparison of arthritic score of non-treated mice (sham), Collagen-induced (CIA) and Liprin-1peptide induced (LIA) mice. Clinical parameters such as slight edema and erythema limited to ankle, slight edema and erythema from the ankle to the tarsal bone, moderate edema and erythema the ankle to the tarsal bone, edema and erythema from the ankle to the entire leg have been considered for clinical score. Statistical significance was determined by 1-way ANOVA: ####: p<0.001 in comparison with the sham group. *Right part:* corresponding cumulative score. (C) Comparison of histological score of non-treated mice (sham), CIA and LIA mice. Statistical significance was determined by 1-way ANOVA: ***: p<0.005; ns: not significant. After 60 days, ankle-joints were evaluated by addressing clinical score from 1 to 4: (1) slight edema and erythema limited to ankle; (2) slight edema and erythema from the ankle to the tarsal bone; (3) moderate edema and erythema the ankle to the tarsal bone; (4): edema and erythema from the ankle to the entire leg. (D) Representative imagines of histological sections of ankle-joints of analyzed animal groups. (E) Effect on pro-inflammatory cytokines profiles (MCP-1, IL-1β, IL-6, IL-12p70, IL-17A, INF-γ, TNF-α,) in sera of treated animals. Statistical significance was determined by 1-way ANOVA: *: p<0.05; ***: p<0.005; ****: p<0.001; ns: not significant. (F) Percentage of CD4+CD25+FoxP3+ Treg cells among CD4+ cells in splenocytes of mice analyzed by flow cytometric analysis. Statistical significance was determined by 1-way ANOVA: **: p<0.01; ns: not significant.

From this point forward, dexamethasone was used as a positive control and administered orally every other day during the treatment phase. As a synthetic glucocorticoid, dexamethasone mediates its anti-inflammatory effects through glucocorticoid receptor–dependent transcriptional regulation, leading to inhibition of NF-κB and AP-1 signaling, suppression of pro-inflammatory cytokine production (including TNF-α, IL-1β and IL-6), and reduced immune cell activation and tissue infiltration^15^.

Consistent with its established mechanism of action, dexamethasone robustly halted the progression of clinical arthritis in this model and served as a benchmark for therapeutic efficacy.

Clinical scoring revealed that all tested administration routes led to a statistically significant reduction in cumulative disease scores in comparison to CIA, with i.v. treatment showing the highest effect, statistically comparable to dexamethasone (Figure 2B, Supplementary Material 1).

Histological analysis at the end of the treatment confirmed the superior performance of i.v. administration in restoring ankle-joints integrity, based on reduced synovial hyperplasia, pannus formation, cellular infiltration and cartilage erosion (Figure 3C-D, Supplementary Material 1). Pro-inflammatory cytokines/chemokine levels in sera were most affected by i.v. and i.p. administrations, with i.v. consistently achieving a higher degree of statistical significance than i.p. In particular, MCP-1 displayed a significant greater reduction than those observed with dexamethasone (Figure 3E, Supplementary Material 1). CD4+CD25+Foxp3+ regulatory T cells (Treg), which play a crucial role in preventing chronic inflammatory and autoimmune diseases by suppressing autoreactive T cells, were significantly increased in splenocyte from CIA mice treated via i.v. injection of TBSV.pLip in comparison to CIA untreated mice. In contrast, i.p. administration resulted in a modest increase in Treg activation, while other routes did not produce statistically significant changes compared to CIA untreated mice (Figure 3F, Supplementary Material 1).

### Induction of arthritis by Liprin-1 peptide

Since RA patients, regardless of RF positivity, produce autoantibodies against the Liprin-1 peptide^16^, we investigated whether Liprin-1 peptide immunization induces arthritis in mice. The dodecamer Liprin-1 peptide (ASVLANVAQAFE) shares 100% homology with the 3 human isoforms Liprin-alpha-1 (NP_803172.1, NP_003617.1, NP_001364935.1) and the 2 mice isoforms (NP_001182015.1, NP_001028491.1). The peptide maps to a central region of the protein that is surface-exposed and annotated as an antigenic region (HPA042271), targeted by a polyclonal antibody.

Male DBA/1J 8-weeks old mice were intradermally injected either with Liprin-1 peptide (0.2 mg/mouse) and chicken type-II Collagen (0.1 mg/mouse) both emulsified in CFA and boosted on day 21 with the same dose (Figure 3A). Within a week post-boost, mice developed progressive inflammatory arthritis (Figure 3B, Supplementary Figure 1). Clinical scores in Liprin1-induced arthritis (LIA) mice were statistically comparable to those in collagen-induced arthritis (CIA) mice throughout the study (Figure 3B, Supplementary Figure 1).

60 days after injection, mice with CIA and LIA showed a significant increase in synovial membrane inflammation, cartilage and bone erosion (Figure 3C-D, Supplementary Figure 1) in comparison to the sham group.

LIA and CIA in mice is associated with a significant increase of most pro-inflammatory cytokines (IL-17, IL-1b, IL-6, IL-12 p60) and of the pro-inflammatory chemokine MCP-1 in sera compared to sham (Figure 3E, Supplementary Figure 1), with IL-17 and MCP-1 being the most abundant response and with a concomitant decrease in the population of Treg cells (Figure 3F, Supplementary Figure 1).

Seven intravenous administrations of 50 µg of TBSV.pLip significantly reduced the progression of clinical arthritis in LIA mice. Treated animals displayed lower cumulative clinical scores compared to controls, and induction of Treg and a general suppression of pro-inflammatory cytokines, indicating effective suppression of disease progression in vivo (Supplementary Figure 2).

### Administration Scheme for Tolerance Induction

Given the unique characteristics of our proteinaceous plant-made nanoparticles in comparison to others previously used in the autoimmune context^17^, we wanted to establish the optimal number of i.v. injections required to effectively halt arthritis progression in the selected animal model by a dose number escalation scheme (Figure 4A).

**Fig. 4.**
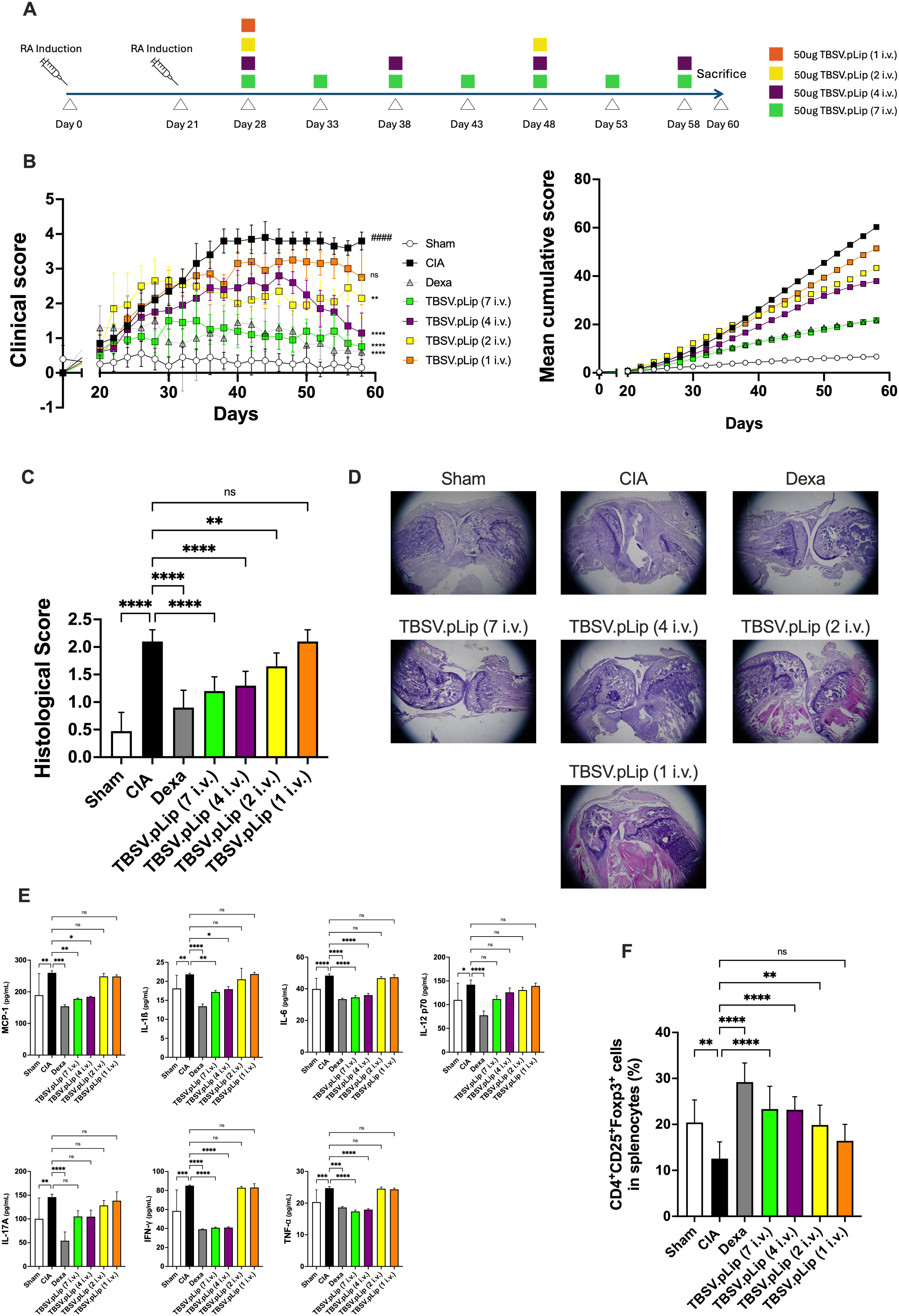
Impact of different dose numbers on TBSV.pLip mediated tolerance. (A) Outline of the experimental design. 8 weeks-old mice received on day 0 and boosted on day 21 intradermally at the basis of the tail 200 µg of collagen with Complete Freund’s adjuvant (CFA). The animals (n=10 for each group) were divided into 7 groups on day 28 based on clinical score and/or paw volume. 50µg of TBSV.pLip were administered i.v. either 7-, 4-, 2- or 1-time i.v. on different days as specified. Control groups included untreated mice (CIA), sham group and a group receiving dexamethasone (1 mg/kg) every other day (dexa). The animals were sacrificed 60 days after CIA induction. (B) *Left part*: Comparison of clinical score of non-treated mice (sham), saline-treated CIA mice (CIA), Dexamethasone-treated CIA mice and mice treated by intravenous administrations of 7, 4, 2 and 1 i.v. doses of TBSV.pLip (50 μg). Clinical parameters such as slight edema and erythema limited to ankle, slight edema and erythema from the ankle to the tarsal bone, moderate edema and erythema the ankle to the tarsal bone, edema and erythema from the ankle to the entire leg have been considered for clinical score. Statistical significance was determined by 1-way ANOVA: **: p<0.01, ****: p<0.001 in comparison with the CIA group; ####: p<0.001 in comparison with the sham group. *Right part:* corresponding cumulative scores. (C) Comparison of histological score in ankle-joints of analyzed animal groups. Statistical significance was determined by 1-way ANOVA: **: p<0.01, ****: p<0.001, ns: not significant, compared with the CIA group. After 58 days, ankle-joints were evaluated by addressing clinical score from 1 to 4: (1) slight edema and erythema limited to ankle; (2) slight edema and erythema from the ankle to the tarsal bone; (3) moderate edema and erythema the ankle to the tarsal bone; (4): edema and erythema from the ankle to the entire leg. (D) Representative imagines of histological sections of ankle-joints of analyzed animal groups. (E) Pro-inflammatory cytokines profiles (MCP-1, IL-1β, IL-6, IL-12p70, IL-17A, INF-γ, TNF-α,) in sera of analyzed mice. Statistical significance was assessed by 1-way ANOVA: *: p<0.05, **: p<0.01; ***: p<0.005; ****: p<0.001. (F) Percentage of CD4+CD25+FoxP3+ Treg cells among CD4+ cells in splenocytes of analyzed mice. Statistical significance was determined by 1-way ANOVA: **: p<0.01, ****: p<0.001, ns: not significant.

Treatments with 2, 4 and 7 i.v. injections of 50 µg TBSV.pLip resulted in a significant reduction in clinical scores compared to the CIA control group (Figure 4B, Supplementary Material 1). These same treatment groups also exhibited markedly lower ankle-joints lesion scores relative to the CIA group (Figure 4 C-D, Supplementary Material 1).

Serum analysis of pro-inflammatory chemokines and cytokines revealed that both the 4 and 7 i.v. TBSV.pLip treatments significantly decreased MCP-1, IL-6, TNF-α, IL-1β and IFN-γ compared to the CIA group (Figure 4E, Supplementary Material 1). The group receiving 1 or 2 i.v. injections of TBSV.pLip showed no reduction in the analyzed cytokines (Figure 4E, Supplementary Material 1). Furthermore, splenocytes analysis demonstrated that mice treated with 4 and 7 i.v. doses of TBSV.pLip restored the percentage of CD4+CD25+Foxp3+ regulatory T cells to levels comparable with healthy controls (Sham Group). A moderate significative increase in this cell population was also observed in the group treated with 2 i.v. injections, relative to the CIA group (Figure 4F, Supplementary Material 1).

Furthermore, the 4- and 7-dose i.v. groups were analyzed at 60 days to assess TBSV.pLip particle accumulation in organs. These groups were selected because they had an identical interval (48 hours) since the last i.v. injection. The data revealed high biological variability and no statistically significant differences among the organs analyzed (Supplementary Figure 3), all of which exhibited low % injected dose per gram of tissue (%ID/g), indicating rapid clearance of the viral particles and minimal accumulation in the examined organs.

### Dose-Effect Relationship of VNPs efficacy

Subsequently, we investigated the dose-response relationship by administering four i.v. doses—5, 20, 50, and 200 µg—every 10 days to assess their ability to induce tolerance. This approach allowed us to evaluate a broader range of concentrations and assess their efficacy in triggering a tolerogenic response. To better approximate the clinical course of RA in humans and to assess preliminary indicators of tolerance durability, we extended the experimental timeline relative to previous studies (Figure 5A).

**Fig. 5.**
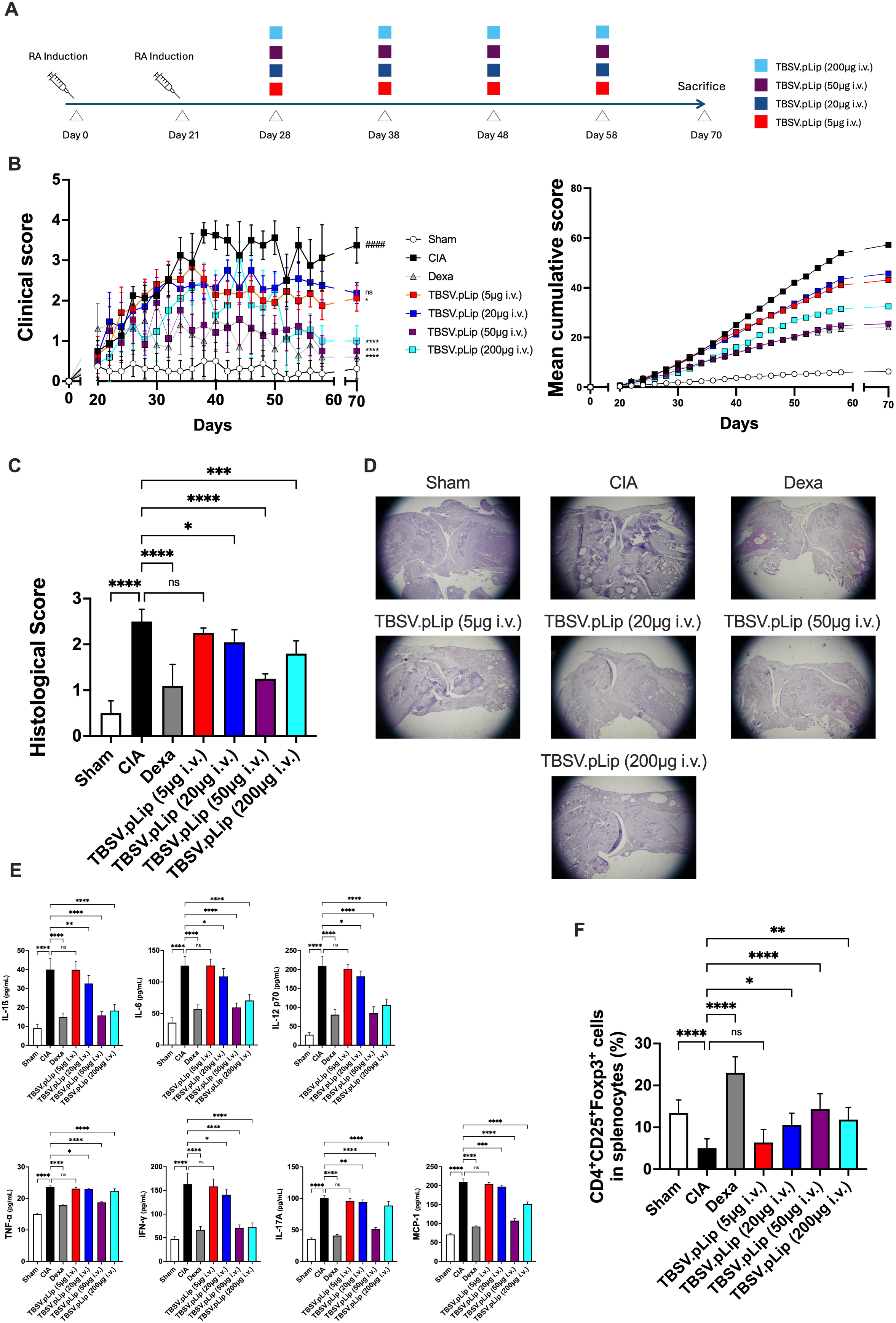
Dose-Effect of TBSV.pLip mediated tolerance. (A) Outline of the experimental design. 8 weeks-old mice received on day 0 and boosted on day 21 intradermally at the basis of the tail 200 µg of collagen with Complete Freund’s adjuvant (CFA). The animals (n=20 for each group) were divided into 7 groups on day 28 based on clinical score and/or paw volume. 5, 20, 50 and 200 µg of TBSV.pLip were administered i.v. 4 times. Control groups included untreated mice (CIA), sham group and a group receiving dexamethasone (1 mg/kg) every other day (Dexa). The animals were sacrificed 70 days after CIA induction. CIA, 3 animals per Sham and TBSV.pLip treated animals were sacrificed for blood collection at days 37, 47 and 57 after CIA induction. (B) *Left part*: Comparison of clinical score of non-treated mice (sham), saline-treated CIA mice (CIA), Dexamethasone-treated CIA mice and mice treated by 4 i.v. administrations of 200, 50, 20, 5 μg of TBSV.pLip. Clinical parameters such as slight edema and erythema limited to ankle, slight edema and erythema from the ankle to the tarsal bone, moderate edema and erythema the ankle to the tarsal bone, edema and erythema from the ankle to the entire leg have been considered for clinical score. Statistical significance was determined by 1-way ANOVA: *: p<0.05, ****: p<0.001, ns: not significant in comparison with the CIA group; ####: p<0.001 in comparison with the sham group. *Right part:* corresponding cumulative scores. (C) Comparison of histological score in ankle-joints of analyzed animal groups. Statistical significance was determined by 1-way ANOVA: *: p<0.05, ***: p<0.005, ****: p<0.001, ns: not significant. (D) Representative imagines of histological sections of ankle-joints of analyzed animal groups. (E) Pro-inflammatory cytokines profiles (MCP-1, IL-1β, IL-6, IL-12p70, IL-17A, INF-γ, TNF-α,) in sera of analyzed mice. Statistical significance was determined by 1-way ANOVA: *: p<0.05, **: p<0.01; ***: p<0.005; ****: p<0.001. (F) Percentage of CD4+CD25+FoxP3+ Treg cells among CD4+ cells in splenocytes of analyzed mice. Statistical significance was determined by 1-way ANOVA: *: p<0.05, **: p<0.01; ***: p<0.005; ****: p<0.001.

Notably, mice receiving four i.v. doses of 50 or 200µg TBSV.pLip exhibited a markedly significant improvement in clinical scores, whereas lower doses (5 and 20µg) failed to produce benefits compared to the CIA control group (Figure 5B, Supplementary Material 1). After 70 days, histological analysis of ankle joints revealed markedly reduced tissue damage in the 50 and 200µg treatment groups relative to CIA controls (Figure 5C-D, Supplementary Material 1).

Correspondingly, serum levels of tested inflammatory cytokines/chemokine were all significantly decreased in mice treated with 50 and 200 µg TBSV.pLip, as well as in the dexamethasone reference group. A moderate but statistically significant reduction was observed in the 20µg group, while the 5µg group showed no significant difference from CIA controls (Figure 5E, Supplementary Material 1).

This anti-inflammatory profile was mirrored in splenic CD4□ CD25□ FoxP3□ regulatory T-cell populations, which were markedly expanded in the 50 and 200µg groups and in dexamethasone-treated mice. The 20µg-group showed partial restoration, whereas no change was detected in the lowest dose group (Figure 5F, Supplementary Material 1).

### Splenectomy does not impact tolerance induction in mice

To elucidate the mechanistic contribution of the spleen in tolerance induction, we compared two groups—splenectomized and non-splenectomized mice—using the experimental protocol corresponding to the four-injection scheme previously used and illustrated in Figure 6A. Splenectomized animals exhibited comparable susceptibility to CIA as control mice, as evidenced by similar clinical scores (Figure 6B, Supplementary Material 1), histopathological assessments (Figure 6C-D, Supplementary Material 1) and serum profiles of most pro-inflammatory cytokines (Figure 6E, Supplementary Material 1), thus confirming the resilience of the model strain to splenectomy, as previously described^18^.

**Fig. 6.**
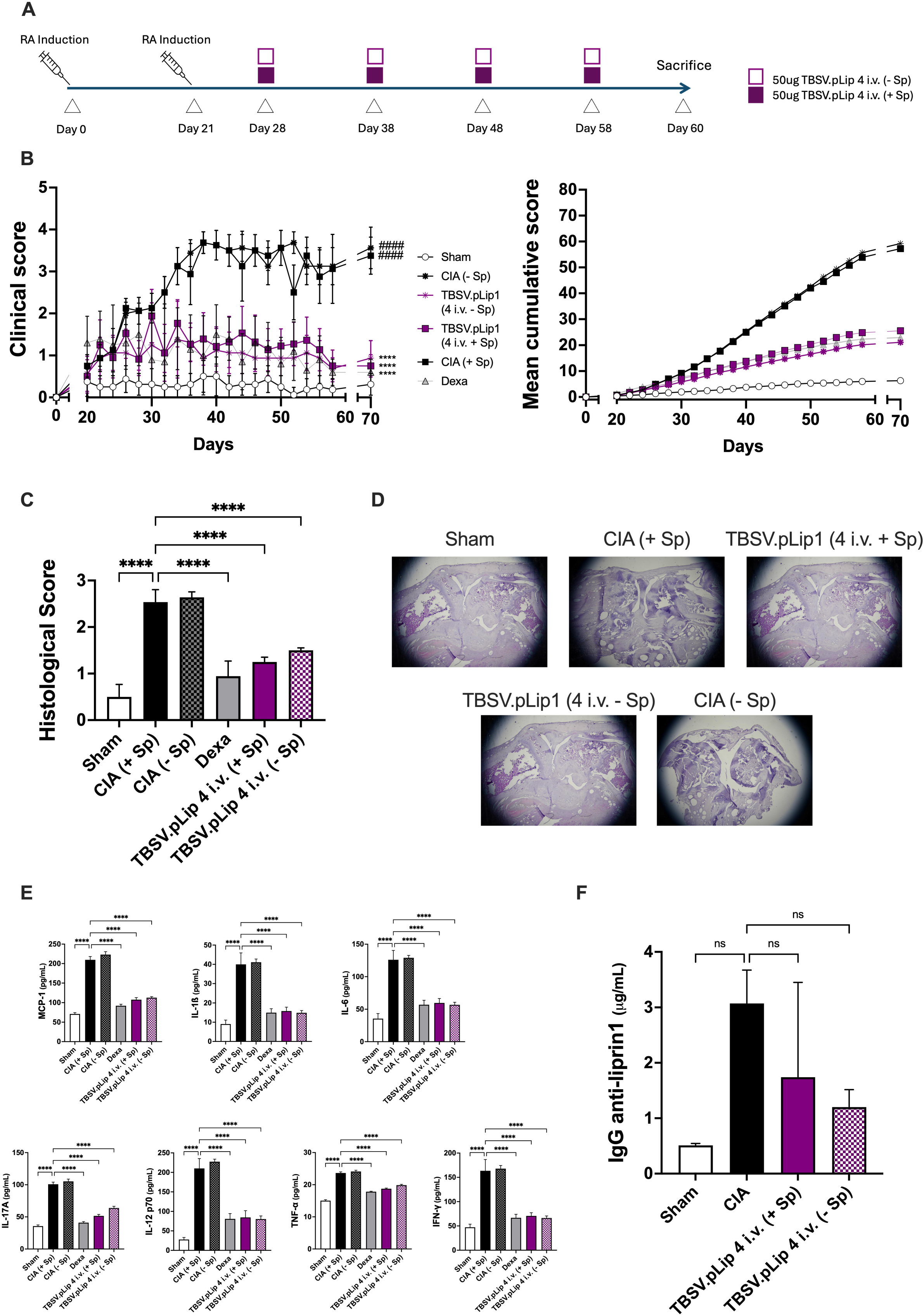
Role of the spleen on the tolerance induction mediated by TBSV.pLip. (A) Outline of the experimental design. 8 weeks-old mice received on day 0 and boosted on day 21 intradermally at the basis of the tail 200 µg of collagen with Complete Freund’s adjuvant (CFA). The animals (n=10 for each group) were divided into 6 groups on day 28 based on clinical score and/or paw volume. 50 µg of TBSV.pLip were administered i.v. 4 times to CIA mice either splenectomized and not. Control groups included untreated mice (CIA) either splenectomized and not, sham group and a group receiving dexamethasone (1 mg/kg) every other day (Dexa). The animals were sacrificed 60 days after CIA induction. (B) *Left part:* Comparison in arthritic score of mice either exposed or not to splenectomy treated by 4 i.v. doses of 50 μg TBSV.pLip and non-treated mice (sham). Clinical parameters such as slight edema and erythema limited to ankle, slight edema and erythema from the ankle to the tarsal bone, moderate edema and erythema the ankle to the tarsal bone, edema and erythema from the ankle to the entire leg have been considered for clinical score. Statistical significance was determined by 1-way ANOVA: *: p<0.05, ****: p<0.001, ns: not significant in comparison with the CIA group; ####: p<0.001 in comparison with the sham group. *Right part:* corresponding cumulative scores. (C) Comparison of histological score in ankle-joints of analyzed animal groups. Statistical significance was determined by 1-way ANOVA: *: p<0.05, **: p<0.01; ***: p<0.005; ****: p<0.001 in comparison to CIA Spleen+. (D) Representative imagines of histological sections of ankle-joints of analyzed animal groups. (E) Pro-inflammatory cytokines profiles (MCP-1, IL-1β, IL-6, IL-12p70, IL-17A, INF-γ, TNF-α,) in sera of analyzed mice. Statistical significance was determined by 1-way ANOVA: *: p<0.05, **: p<0.01; ***: p<0.005; ****: p<0.001. (F) Total IgG in treated mice against the Liprin-1 peptide evaluated by ELISA (n=3 for each group). Statistical significance was determined by 1-way ANOVA: *: p<0.05, **: p<0.01; ***: p<0.005; ****: p<0.001. n.s.: non-significant.

Importantly, mice receiving 4 i.v. doses of 50µg of TBSV.pLip demonstrated a significant reduction in both clinical and histological scores (Figure 6B-D, Supplementary Material 1) regardless of splenectomy status, when compared to untreated CIA controls. Moreover, analysis of serum cytokines revealed a significant decrease in pro-inflammatory mediators in both splenectomized and non-splenectomized groups treated with TBSV.pLip, mirroring the immunosuppressive profile observed in dexamethasone-treated reference animals (Figure 6E, Supplementary Material 1). No antibody reactivity was detected against Liprin-1 peptide in all the groups (Figure 6F).

### Impact of the viral shell on tolerance induction

To indirectly assess whether the observed therapeutic effect was antigen-specific rather than driven by nonspecific properties of the viral scaffold, we compared TBSV.pLip with wild-type TBSV (TBSV WT), which shares an identical structural and physiochemical profile (Figure 1) but does not display the Liprin peptide.

To address this question within a therapeutically relevant setting, we applied the optimized treatment protocol—four i.v. injections of 50 µg VNPs—in CIA mice, directly comparing TBSV.pLip and TBSV WT (Figure 7A). Mice treated with TBSV.pLip showed significant improvement in clinical parameters relative to CIA controls, whereas TBSV WT treatment did not yield any observable benefit (Figure 7B, Supplementary Material 1). Histological analysis corroborated these findings, with reduced joint damage in the TBSV.pLip group and no significant improvement in the TBSV WT group compared to CIA group (Figure 7C–D, Supplementary Material 1).

**Fig. 7.**
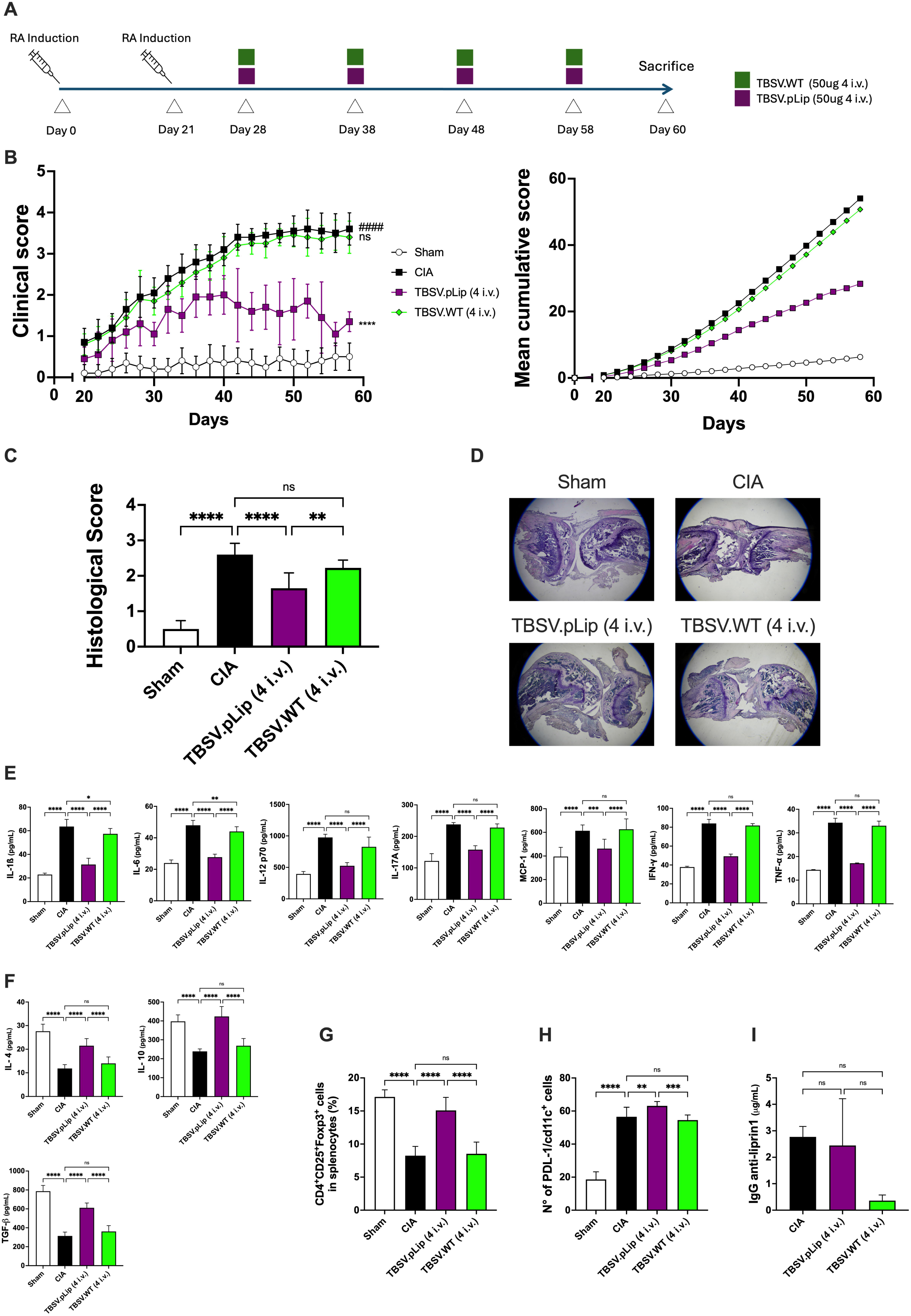
Impact of the viral shell on tolerance induction. (A) Outline of the experimental design. 8 weeks-old mice received on day 0 and boosted on day 21 intradermally at the basis of the tail 200 µg of collagen with Complete Freund’s adjuvant (CFA). The animals (n=10 for each group) were divided into 4 groups on day 28 based on clinical score and/or paw volume. 50 µg of either TBSV.pLip and TBSV WT were administered i.v. 4 times. Control groups included untreated mice (CIA), sham group and a group receiving dexamethasone (1 mg/kg) every other day (Dexa). The animals were sacrificed 60 days after CIA induction. (B) *Left part*: Comparison of clinical score of non-treated mice (sham), saline-treated CIA mice (CIA), Dexamethasone-treated CIA mice and mice treated by 4 intravenous administrations of 50μg of TBSV.pLip and of TBSV WT. Clinical parameters such as slight edema and erythema limited to ankle, slight edema and erythema from the ankle to the tarsal bone, moderate edema and erythema the ankle to the tarsal bone, edema and erythema from the ankle to the entire leg have been considered for clinical score. Statistical significance was determined by 1-way ANOVA: *: p<0.05, ****: p<0.001, ns: not significant in comparison with the CIA group; ####: p<0.001 in comparison with the sham group. *Right part:* corresponding cumulative scores. (C) Comparison of histological score in ankle-joints of analyzed animal groups. Statistical significance was determined by 1-way ANOVA: *: p<0.05, **: p<0.01; ***: p<0.005; ****: p<0.001. (D) Representative imagines of histological sections of ankle-joints of analyzed animal groups. (E) Pro-inflammatory cytokines profiles (MCP-1, IL-1β, IL-6, IL-12p70, IL-17A, INF-γ, TNF-α,) in sera of analyzed mice. Statistical significance was determined by 1-way ANOVA: *: p<0.05, **: p<0.01; ***: p<0.005; ****: p<0.001. (F) Anti-inflammatory cytokines profiles (IL-4, IL-10 and TGF-β) in sera of analyzed mice. Statistical significance was determined by 1-way ANOVA: *: p<0.05, **: p<0.01; ***: p<0.005; ****: p<0.001. (G) Percentage of CD4+CD25+FoxP3+ Treg cells among CD4+ cells in splenocytes of analyzed mice. Statistical significance was determined by 1-way ANOVA: *: p<0.05, **: p<0.01; ***: p<0.005; ****: p<0.001. (H) Immunofluorescence co-labeling analysis of tolerogenic dendritic cells (CD11c□/PD-L1□) in the mesenteric lymph nodes across the experimental groups. *: p<0.05, **: p<0.01; ***: p<0.005; ****: p<0.001. (I) Total IgG in treated mice against the Liprin-1 peptide evaluated by ELISA (n=3 for each group). Statistical significance was determined by 1-way ANOVA: *: p<0.05, **: p<0.01; ***: p<0.005; ****: p<0.001. n.s.: non-significant.

Assessment of pro-inflammatory cytokines revealed overall no significant reduction in the TBSV WT group compared to CIA (except for IL-1β and IL-6), while TBSV.pLip treatment confirmed a robust decrease in all the inflammatory markers considered (Figure 7E, Supplementary Material 1). To explore the anti-inflammatory response, we quantified serum levels of IL-4, IL-10, and TGF-β. TBSV.pLip treatment significantly elevated all three cytokines, with the most pronounced effect observed for TGF-β and IL-4 (Figure 7F, Supplementary Material 1).

Flow cytometric analysis of splenocytes demonstrated a substantial restoration of CD4□ CD25□ Foxp3□ regulatory T cells in the TBSV.pLip-treated group, in contrast to the CIA group and TBSV WT-treated mice, which showed no significant change (Figure 7G, Supplementary Material 1). Immunofluorescence co-labelling of mesenteric lymph nodes revealed a significant increase in tolerogenic dendritic cells (CD11c□/PD-L1□) in the TBSV.pLip group compared to CIA controls. The CIA group exhibited elevated levels of CD11c□/PD-L1□ cells relative to sham, consistent with dendritic cell activation following collagen immunization. Notably, TBSV.pLip treatment further enhanced the presence of tolerogenic dendritic cells, suggesting a role in promoting immune tolerance. In contrast, TBSV WT treatment did not alter dendritic cell populations compared to CIA (Figure 7H, Supplementary Material 1). No differences in antibody reactivity were detected against Liprin-1 peptide in the different groups (Figure 7I).

Given that the two particles, TBSV WT and TBSV.pLip, are structurally equivalent, differing only in the presence of the displayed peptide, these findings support a mechanism driven by the surface-displayed peptide rather than by intrinsic properties of the viral scaffold. The discrepancy with our previous observations, which reported partial immunomodulatory activity of TBSV WT^9^, may be explained by differences in the route of administration (i.p. vs i.v.), known to influence systemic immunomodulation. Additionally, the larger cohort size used in the present study (n = 10 versus n = 3), increases statistical robustness and may account for the clearer discrimination between treatments. Furthermore, the reduced number of administered doses in the current study may have attenuated the effect previously observed in the presence of TBSV WT particles.

## DISCUSSION

Despite substantial advances in disease-modifying therapies for autoimmune disorders, durable restoration of antigen-specific immune tolerance remains an unmet clinical need^19^. Current treatments for rheumatoid arthritis (RA) can achieve remission but require prolonged administration and are associated with systemic immunosuppression. Antigen-specific immunotherapy (ASI) offers the conceptual advantage of selectively targeting pathogenic immune responses while preserving protective immunity^20,21^; however, successful translation to RA has been limited, in part due to the complexity and antigenic diversification of the disease.

In our previous work, administration of a Liprin-derived peptide suppressed disease progression in collagen-induced arthritis (CIA)^9^, despite collagen being the pathogenic antigen driving disease induction. These findings challenged the classical paradigm of strictly antigen-restricted tolerance, suggesting instead that our platform may induce regulatory mechanisms capable of modulating pathogenic immune responses beyond the antigen directly used for tolerization. While antigen restriction represents an advantage in diseases driven by a dominant autoantigen, such as celiac disease^22^, it becomes a limitation in complex autoimmune settings characterized by epitope spreading^23^, such as RA.

Here, we rigorously validated and extended these observations under more stringent experimental conditions. Therapeutic efficacy was reproduced in male DBA/1J mice, which develop a more aggressive CIA phenotype^13^, and cohort size was increased to strengthen statistical robustness. Furthermore, we systematically compared routes of administration and identified intravenous delivery as the most efficacious strategy, consistent with known biodistribution properties of nanoscale materials and their capacity to engage tolerogenic pathways^24,25^.

Importantly, we demonstrate that Liprin-1 itself possesses intrinsic arthritogenic properties, as Liprin-induced inflammatory arthritis recapitulated key clinical and immunological features of CIA in a T cell–dependent manner. These findings establish Liprin-1 as a disease-relevant autoantigen and provide a mechanistic rationale for its use as a therapeutic tolerogen.

Mechanistically, our data support a model in which intravenous TBSV.pLip engages hepatic tolerogenic circuits. The preservation of therapeutic efficacy in splenectomized mice argues against a spleen-dependent mechanism and points toward alternative immunological hubs. The liver is a well-recognized tolerogenic organ in which liver sinusoidal endothelial cells (LSECs) and Kupffer cells promote peripheral tolerance through antigen presentation under sub immunogenic conditions and high expression of inhibitory pathways^26,27^. It is therefore plausible that hepatic antigen capture initiates regulatory programming, leading to FOXP3 Treg expansion, consistent with previous studies demonstrating liver distribution of TBSV particles^28^. The subsequent enrichment of CD11c□ PD-L1□ tolerogenic dendritic cells in secondary lymphoid tissues suggests systemic propagation and stabilization of this regulatory network. PD-L1–expressing dendritic cells are established mediators of peripheral tolerance and contributors to Treg induction and maintenance^29,30^.

Notably, wild-type TBSV particles—despite near-identical physicochemical properties—failed to induce tolerance. The minimal mass difference (∼3.7%) and preserved nanoparticle architecture exclude nonspecific scaffold-driven immunomodulation, generalized immune dampening, or bystander suppression as primary mechanisms. Instead, tolerance induced by TBSV.pLip was strictly dependent on the surface-displayed Liprin-derived peptide, supporting a cargo-driven and antigen-dependent mechanism of immune regulation.

The coordinated increase in circulating IL-4, IL-10, and TGF-β, together with Treg expansion and PD-L1□ tolerogenic dendritic cell induction, is consistent with immune deviation and regulatory reinforcement described in other antigen-specific immunotherapies. IL-4–associated immune deviation has been reported in TCR-engaging immunotherapies such as glatiramer acetate^31,32^ and may cooperate with IL-10 and TGF-β–dependent regulatory circuits to establish durable tolerance.

Collectively, our findings demonstrate that a plant-derived viral nanoparticle platform can induce systemic immune tolerance in experimental arthritis through a peptide-dependent mechanism that transcends strict antigen restriction. This approach offers a potentially scalable and clinically translatable framework for restoring immune homeostasis in complex autoimmune diseases.

## MATERIALS AND METHODS

### Study Design

For animal studies, control and experimental treatments were administered to mice. Sample sizes were determined empirically to ensure adequate statistical power and were consistent with field standards for the techniques used in the study. Prior to treatment initiation, animals were randomized to different treatment groups based on clinical score. The number of samples and the experimental replicates is reported in the figure legends. All procedures involving animals complied with Italian regulations on the protection of animals used for experimental and other scientific purposes (DM 116/1992) as well as EU regulations (Directive legislation (EU) (2010/63/EU).

### Biosynthesis and analysis of nanoparticles

The VNPs were produced and purified as previously described^9^. To confirm the presence, integrity and structure of VNPs total soluble proteins were extracted from infected leaves and analyzed by electrophoresis on agarose gel and DLS, as described^11^. After purification, VNPs were quantified by an ELISA test^11^ and were analyzed manufacturer to assess endotoxin content by LAL test as described by the manufacturer (Gen Script).

Particles size (Z-Average), zeta-potential and polydispersity index (PDI) of VNPs were determined by dynamic light scattering (DLS) and electrophoretic light scattering (ELS) using a Zetasizer Pro-Blue instrument (Malvern-Panalytical, Malvern, UK) and the Malvern Zetasizer software for data analysis. The zeta-potential was calculated using the mixed mode measurement phase analysis light scattering (M3-PALS) technique (Malvern-Panalytical, Malvern, UK). 0.5 µg/µL VNP solutions were dispensed in a folded capillary zeta cell to a final volume of 800 µL using a physiological solution (0.7% NaCl). Data were collected at 25 °C setting the refractive and absorption index to 1.45 and 0.001 respectively. Data are reported as mean ± SD of three technical replicates. All the analyses were performed by Sphera Encapsulation srl, Verona, Italy.

For TEM imaging, samples were absorbed on glow discharged (Leica EM ACE 600) 400 mesh carbon coated copper grids, and then negative stained with 2% aqueous uranyl acetate. Samples were subsequently examined with a Tecnai G2 (FEI) transmission electron microscope operating at 120 kV, and digital images were acquired using a Veleta (Olympus Soft Imaging Solutions) digital camera. Images were analyzed for diameter measurement by ImageJ.

For CP modelling, the primary TBSV and TBSV.pLip coat proteins were modelled independently using Alphafold3^33^. Proteins were aligned using ChimeraX^34^.

### Mice

Male DBA/1J mice (8 weeks; Envigo, Italy) were used for these studies. Mice were housed in individual cages (two for each group) and maintained under a 12:12 light–dark cycle at 21°C ± 1°C and 50% ± 5% humidity. The animals were acclimated to their environment for 1 week and had ad libitum access to tap water and a standard rodent diet.

### Induction of Arthritis

Induction of arthritis was performed by inoculation with Chicken type-II collagen with Complete Freund’s adjuvant (CFA) on Day 0 (0.1 mg/0.1 mL/mouse), intradermally (ID) at the basis of the tail and boosted ID on day 21 with collagen emulsified with IFA/CFA (TBD) (0.1 mg/0.1 mL/mouse)^35^. The positive control with dexamethasone was administered orally.

### Clinical and histological assessment of arthritis

Mice were assessed two times per week for redness and swelling of limbs and a clinical score from 1 to 4 (i) slight edema and erythema limited to ankle; (ii) slight edema and erythema from the ankle to the tarsal bone; (iii) moderate edema and erythema the ankle to the tarsal bone; (iv): edema and erythema from the ankle to the entire leg)^36^. At the end of the experiment, the paws from the mice were collected and prepared for hematoxylin and eosin staining (H&E) and evaluated for histological alterations. Moreover, at the end of experiment, spleen weights for all the groups were recorded.

### FACS Analysis

Spleen were collected from mice, washed twice with phosphate-buffered saline (PBS), ground with two glass slides, filtered with a 200-mesh sieve, and centrifuged at 1000 rpm for 5 min; the supernatant was removed; and the precipitate was. The precipitate was held twice with PBS, centrifuged at 1000 rpm for 10 min; the supernatant was removed; and the precipitate was left. Then, 2 mL of erythrocyte lysate was added to resuspend the precipitate, incubated at room temperature for 5 min, and centrifuged at 1000 rpm for 10 min to obtain splenocytes. A single-cell suspension of splenocytes at 2×106/mL was prepared. Cells were stained with anti-CD4, anti-CD25, and anti-Foxp3 monoclonal antibodies according to the instructions of the manufacturer. Cell events were collected on a flow cytometer and the stained CD4+CD25+Foxp3+Treg cells were analyzed using flow software^37^.

### Cytokine Quantification

Serum was collected and used for cytokines analysis (IL-12 p70, IL-1b, INF-_γ_, IL-6, MCP-1, IL17 A and TNF-_α_) by performing ELISA kit and following manufacturer’s instructions.

### ELISA for detecting anti-Liprin-1 antibodies

ELISA plates were coated with 100µl of 10µg/ml of streptavidin in carbonate buffer at pH 9.6. A standard curve was made with purified mouse IgG. After plate blocking, 1 µg/ml of biotin-Lip1 was added and incubated overnight at 4°C. Plates were washed and 1:1 serial dilutions of animal sera were added. Plates were washed and a 1:5000 dilution of anti-mouse AP- secondary antibody was added in PBS 1X +1%BSA and incubated 1 hour at 25°C. Plates were then washed and developed with pNPP 1h at 37°C.

### Splenectomy

The splenectomy was performed as previously described^38^. Animals were anesthetized with 5% isoflurane and monitored until their respiratory rate dropped below 60 breaths per minute. Surgical instruments were sterilized between procedures and arranged on a sterile table along with sutures, forceps, scissors, and twisters. The abdominal skin was sterilized with iodine, and a 0.5 cm midline incision was made just below the xyphoid process. The underlying fascia was identified and incised along the same length. The spleen was mobilized from the left upper quadrant using smooth forceps and a cotton-tip applicator and brought to the midline incision. Attachments and vessels at the hilum were cauterized using an eye cautery with a heated tip, applying a vessel-preserving technique to minimize trauma and blood loss. The fascial layer was closed with 4-0 Vicryl sutures, and the skin was closed with five interrupted 4-0 Vicryl stitches using a Wire Twister. Procedures lasted approximately 5–10 minutes. Animals were kept under a heat lamp to prevent hypothermia and received 1 mL of subcutaneous saline to replace fluid losses. Postoperatively, mice were observed for several minutes and then hourly for 4 hours.

### Preparation of Mouse Organs for TBSV.pLip Particle Content Analysis

Tissue samples were placed in 1.5 mL microcentrifuge tubes containing three glass beads. An extraction buffer (25 mM bicine, 150 mM NaCl, 1% Triton X-100, pH 7.6) was added at a ratio of 1:2 (w/v). Samples were homogenized for 6 seconds at maximum speed, followed by incubation on ice for 20 minutes. The homogenate was centrifuged at 5,000 × g for 10 minutes at 4 °C. The resulting supernatant was carefully collected and transferred to a new 1.5 mL microcentrifuge tube for ELISA analysis.

### Statistical Analysis

All values are expressed as mean ± standard deviation (SD) of the mean of n observations. Data sets were examined by one-way analysis of variance ANOVA followed by a Bonferroni post-hoc test for multiple comparisons. Data of DLS are presented as mean +/- standard deviation. Statistical significance between groups was determined using a two-tailed Student’s t-test. A p-value< 0.05 was considered statistically significant.

## List of Supplementary Materials

**Table S1:** Data file reporting all the statistical analysis performed for all pre-clinical experiments reported.

**Figure S2: Tolerance induction in LIA mice by TBSV.pLip**

(A) Outline of the experimental design. 8 weeks-old mice received on day 0 and boosted on day 21 intradermally at the basis of the tail 200 µg of Liprin-1 peptide with Complete Freund’s adjuvant (CFA). The animals (n=10 for each group) were divided into 4 groups on day 28 based on clinical score and/or paw volume. 50 µg of TBSV.pLip were administered i.v. Control group of untreated mice (sham group). The animals were sacrificed 60 days after LIA induction.

(B) *Left part*: Comparison of clinical score of non-treated mice (sham), Liprin-1 peptide-treated mice (LIA) and mice treated by intravenous administrations of 50µg of TBSV.pLip. Clinical parameters such as slight edema and erythema limited to ankle, slight edema and erythema from the ankle to the tarsal bone, moderate edema and erythema the ankle to the tarsal bone, edema and erythema from the ankle to the entire leg have been considered for clinical score. Statistical significance was determined by 1-way ANOVA: *: p<0.05, ****: p<0.001, ns: not significant in comparison with the CIA group; ####: p<0.001 in comparison with the sham group. *Right part:* corresponding cumulative scores.

(C) Comparison of histological score in ankle-joints of analyzed animal groups. Statistical significance was determined by 1-way ANOVA: *: p<0.05, **: p<0.01; ***: p<0.005; ****: p<0.001.

(D) Representative imagines of histological sections of ankle-joints of analyzed animal groups.

(E) Pro-inflammatory cytokines profiles (MCP-1, IL-1β, IL-6, IL-12p70, IL-17A, INF-γ, TNF-α,) in sera of analyzed mice. Statistical significance was determined by 1-way ANOVA: *: p<0.05, **: p<0.01; ***: p<0.005; ****: p<0.001.

(F) Percentage of CD4+CD25+FoxP3+ Treg cells among CD4+ cells in splenocytes of analyzed mice. Statistical significance was determined by 1-way ANOVA: *: p<0.05, **: p<0.01; ***: p<0.005; ****: p<0.001.

**Figure S3: Biodistribution of TBSV.pLip following intravenous administration.**

Tissue uptake is expressed as percentage of injected dose per gram of tissue (%ID/g) in major organs (liver, spleen, kidney, stomach, heart, lungs, and brain). Purple bars represent TBSV.pLip 4 intravenous injections (4 i.v.), while green bars represent 7 intravenous injections (7 i.v.). Data are shown as mean ± standard deviation. In purple: 7 i.v. injections of TBSV.pLip, in green: 4 i.v. injections of TBSV.pLip.

**Figure S4: Assessment of combined i.v. and s.c. administrations of TBSV.pLip for Tolerance Induction**

(A) Outline of the experimental design. 8 weeks-old mice received on day 0 and boosted on day 21 intradermally at the basis of the tail 200 µg of collagen with Complete Freund’s adjuvant (CFA). The animals (n=10 for each group) were divided into 4 groups on day 28 based on clinical score and/or paw volume. 50 µg of TBSV.pLip were administered i.v. and/or s.c. as indicated. Control groups included untreated mice (CIA), sham group and a group receiving dexamethasone (1 mg/kg) every other day (Dexa). The animals were sacrificed 60 days after CIA induction.

(B) *Left part*: Comparison of clinical score of non-treated mice (sham), saline-treated CIA mice (CIA), Dexamethasone-treated CIA mice and mice treated by sub-cutaneous and/or intravenous administrations of 50µg of TBSV.pLip. Clinical parameters such as slight edema and erythema limited to ankle, slight edema and erythema from the ankle to the tarsal bone, moderate edema and erythema the ankle to the tarsal bone, edema and erythema from the ankle to the entire leg have been considered for clinical score. Statistical significance was determined by 1-way ANOVA: *: p<0.05, ****: p<0.001, ns: not significant in comparison with the CIA group; ####: p<0.001 in comparison with the sham group. *Right part:* corresponding cumulative score.

(C) Comparison of histological score in ankle-joints of analyzed animal groups. Statistical significance was determined by 1-way ANOVA: *: p<0.05, **: p<0.01; ***: p<0.005; ****: p<0.001.

(D) Representative imagines of histological sections of ankle-joints of analyzed animal groups.

(E) Pro-inflammatory cytokines profiles (MCP-1, IL-1β, IL-6, IL-12p70, IL-17A, INF-γ, TNF-α,) in sera of analyzed mice. Statistical significance was determined by 1-way ANOVA: *: p<0.05, **: p<0.01; ***: p<0.005; ****: p<0.001.

(F) Percentage of CD4+CD25+FoxP3+ Treg cells among CD4+ cells in splenocytes of analyzed mice. Statistical significance was determined by 1-way ANOVA: *: p<0.05, **: p<0.01; ***: p<0.005; ****: p<0.001.

## ACKNOWLEDGMENTS

Authors acknowledge CPT (Centro Piattaforme Tecnologiche) of the University of Verona for the access and support to the DLS equipment.

## AUTHOR CONTRIBUTIONS

ARo, ARa, EG, and RZ contributed to the experimental design related to particle production. MC and EE performed and analyzed the animal model experiments. JEB, JM, NS, EB, RZ, and LA contributed to the overall experimental design of the study. AmC contributed to the analysis of IgG responses and established and performed the ELISA assays. AR performed the statistical analyses. LA coordinated the study, secured funding, and wrote the manuscript. All authors reviewed and approved the final version of the manuscript.

## DATA AND MATERIAL AVAILABILITY

All raw data relative to the experiments reported are present in Table 1 of the Supplementary Materials

