## Supplementary figures and images for "Targeting autoimmunity in Rheumatoid Arthritis with nanoparticles displaying Liprin-1 peptide"

### SI_Figura S2 copia.tiff

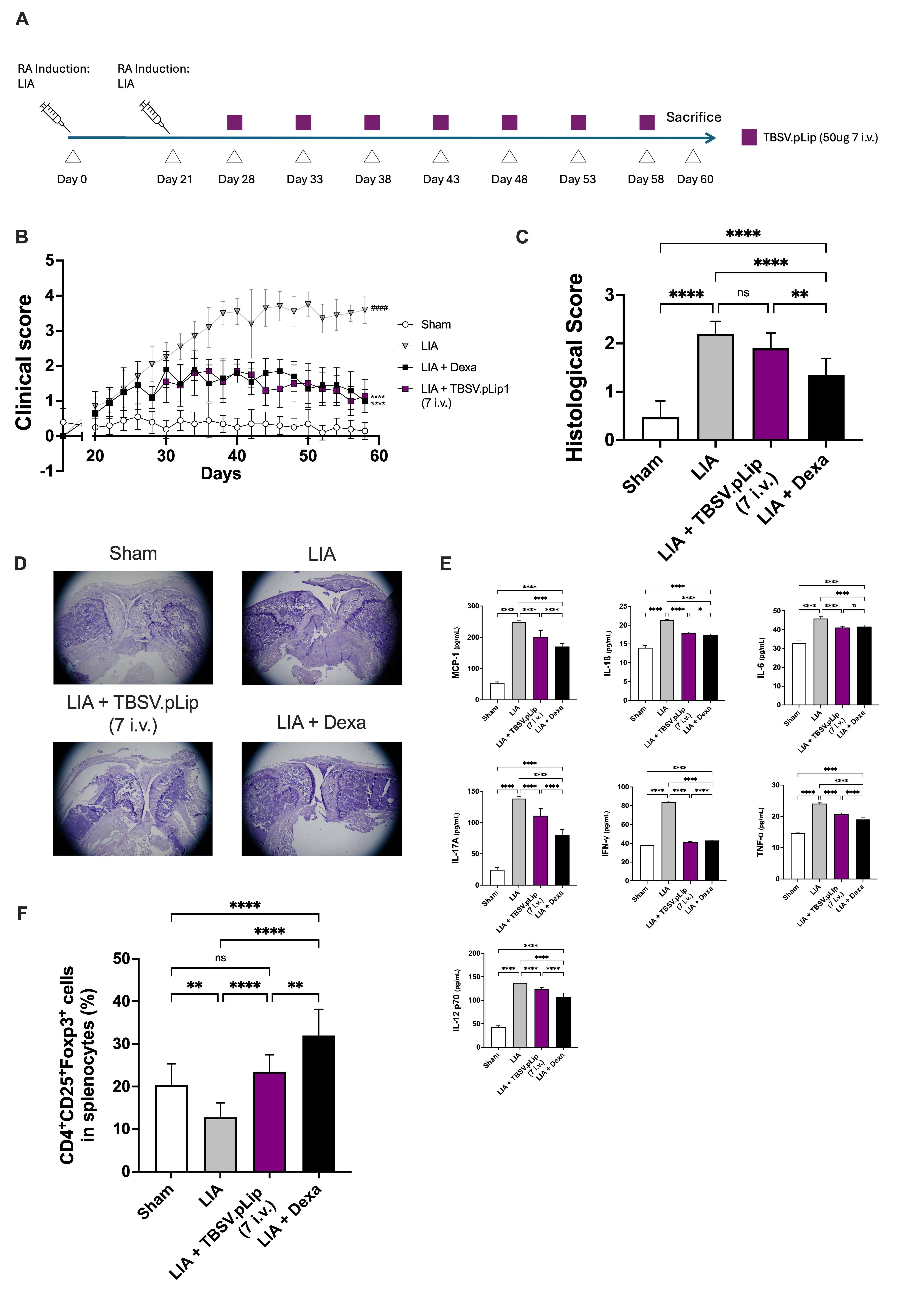

### SI_Figura S4 copia.tiff

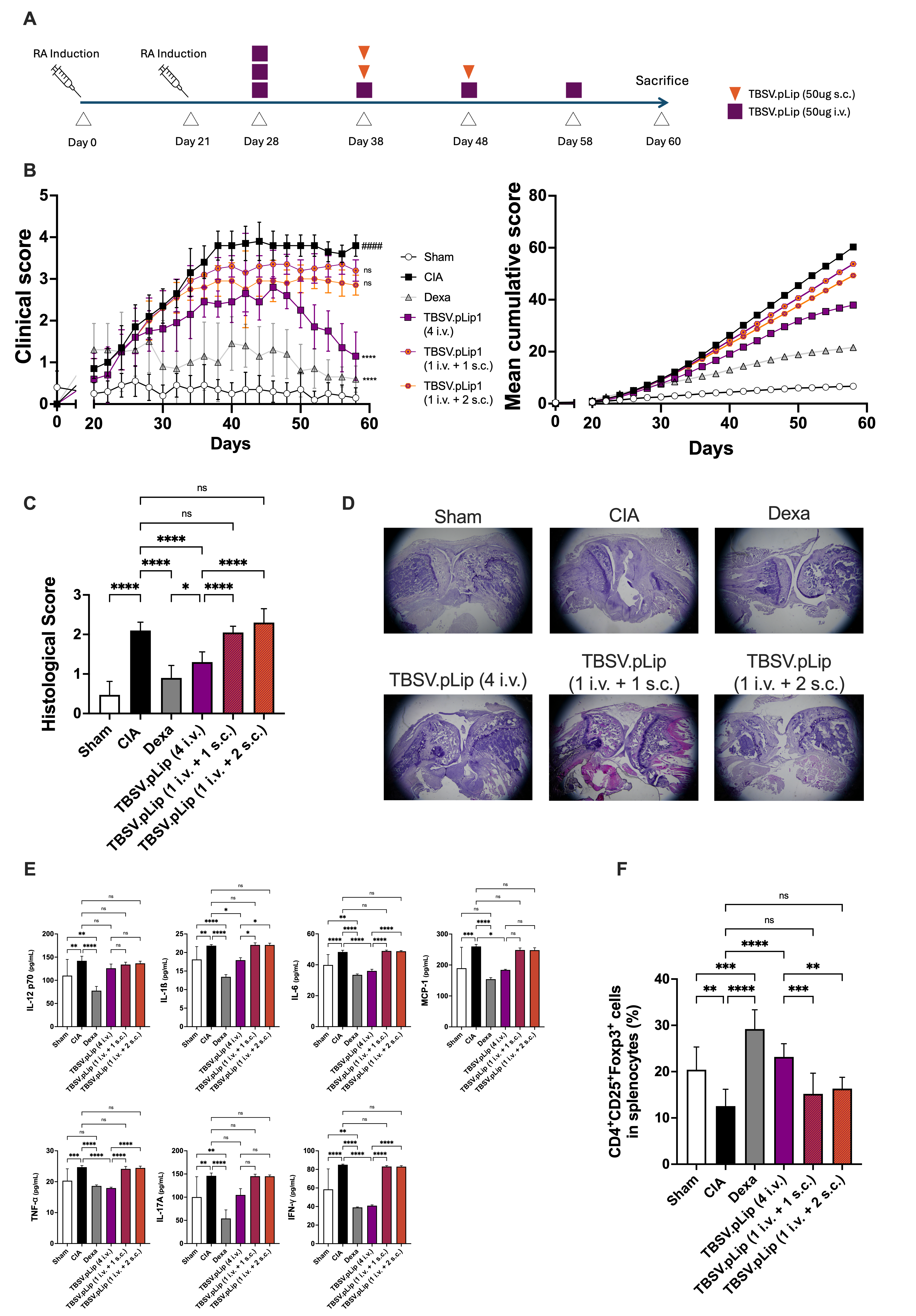

### Supplementary Figure 3

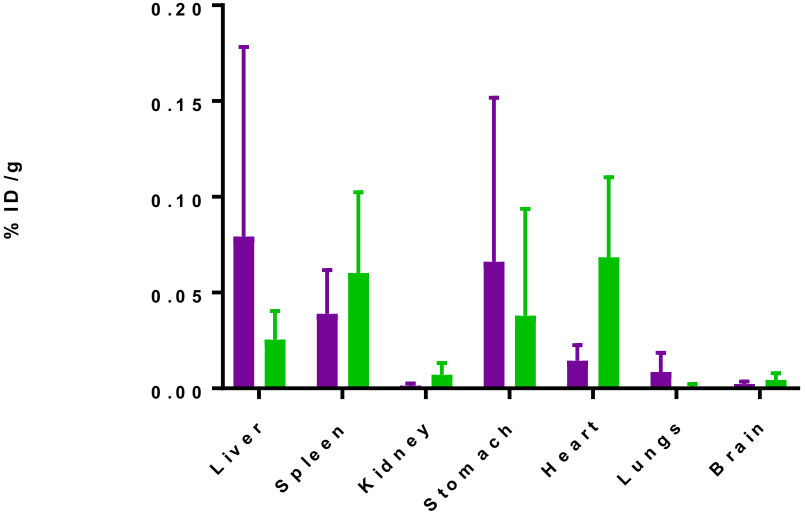
